# Feasibility of FreeSurfer processing for T1-weighted brain images of 5-year-olds: semiautomated protocol of FinnBrain Neuroimaging Lab

**DOI:** 10.1101/2021.05.25.445419

**Authors:** Elmo P. Pulli, Eero Silver, Venla Kumpulainen, Anni Copeland, Harri Merisaari, Jani Saunavaara, Riitta Parkkola, Tuire Lähdesmäki, Ekaterina Saukko, Saara Nolvi, Eeva-Leena Kataja, Riikka Korja, Linnea Karlsson, Hasse Karlsson, Jetro J. Tuulari

**Author notes:** Corresponding author, Elmo P. Pulli, FinnBrain Birth Cohort Study, Teutori building, 2^nd^ floor, Lemminkäisenkatu 3, 21520, Turku, Finland.

## Abstract

Pediatric neuroimaging is a quickly developing field that still faces important methodological challenges. One key challenge is the use of many different atlases, automated segmentation tools, manual edits in semiautomated protocols, and quality control protocols, which complicates comparisons between studies. In this article, we present our semiautomated segmentation protocol using FreeSurfer v6.0, ENIGMA consortium software, and the quality control protocol that was used in FinnBrain Birth Cohort Study. We used a dichotomous quality rating scale for inclusion and exclusion of images, and then explored the quality on a region of interest level to exclude all regions with major segmentation errors. The effects of manual edits on cortical thickness values were minor: less than 2% in all regions. Supplementary materials cover registration and additional edit options in FreeSurfer and comparison to the computational anatomy toolbox (CAT12). Overall, we conclude that despite minor imperfections FreeSurfer can be reliably used to segment cortical metrics from T1-weighted images of 5-year-old children with appropriate quality assessment in place. However, custom templates may be needed to optimize the results for the subcortical areas. Our semiautomated segmentation protocol provides high quality pediatric neuroimaging data and could help investigators working with similar data sets.

## Introduction

There are multiple methodological challenges in pediatric neuroimaging studies that may affect quality of data and comparisons between studies. Magnetic resonance imaging (MRI) requires the subject to lie still while awake, which is more of a challenge with children than with adults (Blumenthal et al., 2002; Poldrack et al., 2002; Theys et al., 2014). One study (Blumenthal et al., 2002) found that mild, moderate, and severe motion artefact were associated with 4%, 7%, and 27% loss of total gray matter (GM) volume in segmentation, respectively. Furthermore, subtle motion can cause bias even when visible artefact is absent (Alexander-Bloch et al., 2016). Another core challenge is the variation in preprocessing and segmentation techniques (Phan, Smeets, et al., 2018), due to a lack of a “gold standard” processing pipeline for pediatric brain images. Some studies develop their own pediatric atlases (Fonov et al., 2011; Kuklisova-Murgasova et al., 2011; Oishi et al., 2011), while others apply segmentation tools developed for adult imaging that may possess limitations when applied to pediatric imaging data (Phan, Sima, et al., 2018; Wilke et al., 2003; Yoon et al., 2009). Therefore, some studies rightfully emphasize the importance of a validated quality control protocol (Schoemaker et al., 2016). Other potential challenges when utilizing software with adult templates in pediatric imaging poorer contrast between GM and white matter (WM) as well as significant differences in brain volumes (Schoemaker et al., 2016). These challenges are most pronounced in the first years of life, hence the development of specialized pipelines for this age group such as infant FreeSurfer for 0- to 2-year-olds (Zöllei et al., 2020).

FreeSurfer (http://surfer.nmr.mgh.harvard.edu/) is an open source software suite for processing brain MRI images that is commonly used in pediatric neuroimaging (Al Harrach et al., 2019; Barnes-Davis et al., 2020; Black et al., 2012; Boutzoukas et al., 2020; Clark et al., 2014; El Marroun et al., 2016; Garnett et al., 2018; Ghosh et al., 2010; Lee et al., 2017; Nwosu et al., 2018; Phan, Smeets, et al., 2018; Ranger et al., 2013; Roos et al., 2014; Wedderburn et al., 2020). The automated FreeSurfer segmentation protocol utilizes surface-based parcellation of cortical regions based on cortical folding patterns and a priori knowledge of anatomical structures (further technical information in Dale et al., 1999; Fischl, Sereno, & Dale, 1999). The FreeSurfer instructions recommend to visually check and, when necessary, manually edit the data. The manual edits can fix errors in the automated segmentation such as skull-stripping, WM, or pial errors (errors in the outer border of cortical GM). The FreeSurfer instructions suggest that this process takes approximately 30 minutes. However, in our experience, this timeframe seems far too short for careful quality assessment and editing. The time requirement is perhaps the most important challenge in manual editing of brain images. Another one is the fact that the edits may lead to inter- and intra-rater bias. Nevertheless, effects of motion artefact must be considered in the segmentation process (Blumenthal et al., 2002), as some systematic errors in pial border, subcortical structures, and the cerebellum have been observed in structural brain images of 5-year-olds without manual edits (Phan, Smeets, et al., 2018). While a visual check for major errors has obvious benefits, the benefits of manual edits are not as clear in children (Beelen et al., 2020) or adults (Guenette et al., 2018; McCarthy et al., 2015; Waters et al., 2019) as errors that can be manually edited are often small and therefore only have minor effects on cortical thickness (CT) values. Consequently, they do not necessarily affect the significant findings in group comparisons (McCarthy et al., 2015) or brain-behavior relationships (Waters et al., 2019). However, we argue that systematic manual edits can help with quality control as they simultaneously maximize the chance to find segmentation errors that can be subsequently fixed.

Quality control is often done by applying a dichotomous pass or fail scale: either by simply excluding the cases with excessive motion artefact (Boutzoukas et al., 2020; Garnett et al., 2018; Ranger et al., 2013; Vanderauwera et al., 2018; Yang et al., 2016; Yang et al., 2015), excluding issues related to pathologies (Al Harrach et al., 2019; Ranger et al., 2013), excluding extreme outlier cases (Nwosu et al., 2018), or it is simply noting that all images were considered to be of sufficient quality without a more detailed description of the criteria (Barnes-Davis et al., 2020). Another approach is to rate the image on a Likert scale from excellent or no motion artefact to unusable (Blumenthal et al., 2002; White et al., 2018). One key challenge with this approach that the exact borders between categories are very difficult to describe accurately in writing, and terms such as “subtle” and “significant” concentric bands or motion artefact are frequently used to draw the borders (Blumenthal et al., 2002; Shaw et al., 2007). Consequently, even if good intra- and inter-rater reliability can be reached within a study (Shaw et al., 2007), there can be large differences in how different studies define the categories. In many cases, the line of exclusion is drawn between moderate and severe (Lyall et al., 2015) or mild and moderate artefact (Shaw et al., 2007), and either way this fundamentally results in two categories: images with acceptable quality and images with unacceptable quality. Instead of a further quality classification via a Likert scale based on the amount of visible artefact, it might be beneficial to quality check all regions of interest (ROI) separately to verify high quality of the data. Especially considering the fact that the developing brain undergoes multiple non-linear growth patterns (Phan, Smeets, et al., 2018; Wilke et al., 2003), which may cause issues when utilizing an adult template (Muzik et al., 2000; Phan, Sima, et al., 2018; Yoon et al., 2009), and local errors related to this challenge may be missed if quality check is based solely on the severity of visible motion artefact.

In this article, we propose a dichotomous rating scale for inclusion and exclusion of images, combined with a post-processing quality control protocol to visually confirm high quality data on a ROI level. For the automated segmentation tool in this protocol, we chose FreeSurfer based on the following practical advantages: 1) FreeSurfer has been validated for use in children between ages four and eleven years (Ghosh et al., 2010), and multiple studies have used FreeSurfer to find brain associations between brain structure and risk factors or cognitive differences in children (Black et al., 2012; Clark et al., 2014; Wedderburn et al., 2020); 2) FreeSurfer provides a method to accurately assess image quality and to fix certain types of errors via Freeview; and 3) Rigorous quality control protocols, such as the one provided by the ENIGMA consortium (Enhancing Neuro Imaging Genetics through Meta Analysis; http://enigma.ini.usc.edu/), already exist for FreeSurfer to make final quality assessment on such a level that allows the researchers to exclude single ROIs with imperfect segmentation. We decided to use the ENIGMA quality control protocol, as it is widely used and accepted (Thompson et al., 2020), and has been successfully implemented for both adults (Thompson et al., 2020) and children (Boedhoe et al., 2018; Hoogman et al., 2019). The manual edits instructed by FreeSurfer and this rigorous ENIGMA quality control protocol were combined to create the semiautomated segmentation pipeline used in the FinnBrain Neuroimaging Lab.

In the current study, we used a representative subsample of ca. 5-year-olds that participated in MRI brain scans as part of the FinnBrain Birth Cohort Study. We give a detailed description of our manual editing and quality control protocol for T1-weighted MRI images in the FreeSurfer software suite. In addition, we used the ENIGMA quality control protocol and compare the findings to our protocol. Furthermore, in a complementary analysis, we compared automated segmentation results between FreeSurfer and the statistical parametric mapping (SPM; https://fil.ion.ucl.ac.uk/spm) based computational anatomy toolbox (CAT12; http://www.neuro.uni-jena.de/cat/) to assess to the level of agreement. Finally, we compared the standard recon-all to other optional flags in FreeSurfer.

## Methods

This study was conducted in accordance with the Declaration of Helsinki, and it was approved by the Joint Ethics Committee of the University of Turku and the Hospital District of Southwest Finland (07.08.2018) §330, ETMK: 31/180/2011.

### Participants

The participants are part of the FinnBrain Birth Cohort Study (www.finnbrain.fi) (Karlsson et al., 2018), where 5-year-olds were invited to neuropsychological, logopedic, neuroimaging, and pediatric study visits. For the neuroimaging visit, we primarily recruited participants that had a prior visit to neuropsychological measurements at ca. 5 years of age (n = 141/146). However, there were a few exceptions: three participants were included without a neuropsychological visit, as they had an exposure to maternal prenatal synthetic glucocorticoid treatment (recruited separately for a nested case–control sub-study). The data additionally includes two participants that were enrolled for pilot scans. We aimed to scan all subjects between the ages 5 years 3 months and 5 years 5 months, and 135/146 (92%) of the participants attended the visit within this timeframe. The exclusion criteria for this study were: 1) born before gestational week 35 (before gestational week 32 for those with exposure to maternal prenatal synthetic glucocorticoid treatment), 2) developmental anomaly or abnormalities in senses or communication (e.g. blindness, deafness, congenital heart disease), 3) known long-term medical diagnosis (e.g. epilepsy, autism), 4) ongoing medical examinations or clinical follow up in a hospital (meaning there has been a referral from primary care setting to special health care), 5) child use of continuous, daily medication (including per oral medications, topical creams and inhalants. One exception to this was desmopressin (®Minirin) medication, which was allowed), 6) history of head trauma (defined as concussion necessitating clinical follow up in a health care setting or worse), 7) metallic (golden) ear tubes (to assure good-quality scans), and routine MRI contraindications.

In the current study, we used a subsample (approximately two thirds of the full sample) that consists of the participants that were scanned before a temporary stop to visits due to the restrictions caused by the COVID-19 pandemic. The scans were performed between 29 October 2017 and 1 March 2020. We contacted 415 families and reached 363 (87%) of them. In total, 146 (40% of the reached families) participants attended imaging visits (one pair of twins, one participant attended twice). Eight of them did not start the scan, and four were excluded due to excess motion artefact in the T1-image. Thereafter, 134 T1 images entered the processing pipelines. Supplementary Table 1 presents the demographic data as recommended in our earlier review (Pulli et al., 2019).

### The study visits

All MRI scans were performed for research purposes by the research staff (one research nurse, four PhD students, and two MR technologists). Before the visit, each family was personally contacted and recruited via telephone calls by a research staff member. The scan preparations started with the recruitment and at home training. We introduced the image acquisition process to the parents and advised them to explain the process to their children and confirm child assent before the follow up phone call that was used to confirm the willingness to participate. Thereafter, we advised the parents to use at home familiarization methods such as showing a video describing the visit, playing audio of scanner sounds, encouraging the child to lie still like a statue (“statue game”), and practicing with a homemade mock scanner, e.g., a cardboard box with a hole to view a movie through. The visit was marketed to the participants as a “space adventure”, which is in principle similar to the previously described “submarine protocol” (Theys et al., 2014) but the child was allowed to come up with other settings as well. A member of the research staff made a home visit before the scan to deliver earplugs and headphones, to give more detailed information about the visit, and to answer any remaining questions. An added benefit of the home visit was the chance to meet the participating child and that way start the familiarization with the research staff.

At the start of the visit, we familiarized the participant with the research team (research nurse and a medically trained PhD student) and acquired written informed consent from both parents. This first portion of the visit included a practice session using a non-commercial mock scanner consisting of a toy tunnel and a homemade wooden head coil. Inexpensive non-commercial mock scanners have been shown to be as effective as commercial ones (Barnea-Goraly et al., 2014). The participants brought at least one of their toys that would undergo a mock scan (e.g., an MRI compatible stuffed animal they could also bring with them into the real scanner). The researcher played scanner sounds on their cell phone during the mock scan and the child could take pictures of the toy lying still and of the toy being moved by the researcher to demonstrate the importance of lying still during the scan. Communication during the scan was practiced. Overall, these preparations at the scan site were highly variable as we did our best to accommodate to befit the child characteristics (e.g., taking into account the physical activity and anxiety) in cooperation with the family. Finally, we served a light meal of the participant’s choice before the scan.

The participants were scanned awake or during natural sleep. One member of the research staff and parent(s) stayed in the scanner room throughout the whole scan. During the scan, participants wore earplugs and headphones. Through the headphones, they were able to listen to the movie or TV show of their choice while watching it with the help of mirrors fitted into the head coil (the TV was located at the foot of the bed of the scanner). Some foam padding was applied to help the head stay still and assure comfortable position. Participants were given a “signal ball” to throw in case they needed or wanted to stop or pause the scan (e.g., to visit the toilet). If the research staff member noticed movement, they gently reminded the participant to stay still by lightly touching their foot. This method of communication was agreed on earlier in the visit and was planned to convey a clear signal of presence while minimizing the tactile stimulation. Many of the methods used to reduce anxiety and motion during the scan have been described in earlier studies (Epstein et al., 2007; Greene et al., 2016).

All images were viewed by one neuroradiologist (RP) who then consulted a pediatric neurologist (TL) when necessary. There were four (out of 146, 2.7%) cases with an incidental finding that required consultation. The protocol with incidental findings has been described in our earlier work (Kumpulainen et al., 2020), and a separate report of their incidence is in preparation for the eventual full data set.

### MRI data acquisition

Participants were scanned using a Siemens Magnetom Skyra fit 3T with a 20-element head/neck matrix coil. We used Generalized Autocalibrating Partially Parallel Acquisition (GRAPPA) technique to accelerate image acquisition (parallel acquisition technique [PAT] factor of 2 was used). The MRI data was acquired as a part of max. 60-minute scan protocol. The scans included a high resolution T1 magnetization prepared rapid gradient echo (MPRAGE), a T2 turbo spin echo (TSE), a 7-minute resting state functional MRI, and a 96-direction single shell (b = 1000 s/mm^2^) diffusion tensor imaging (DTI) sequence (Merisaari et al., 2019) as well as a 31-direction with b = 650 s/mm^2^ and a 80-direction with b = 2000 s/mm^2^. For the purposes of the current study, we acquired high resolution T1-weighted images with the following sequence parameters: TR = 1900 ms, TE = 3.26 ms, TI = 900 ms, flip angle = 9 degrees, voxel size = 1.0 × 1.0 × 1.0 mm^3^, FOV 256 × 256 mm^2^. The scans were planned as per recommendations of the FreeSurfer developers (https://surfer.nmr.mgh.harvard.edu/fswiki/FreeSurferWiki?action=AttachFile&do=get&target=FreeSurfer_Suggested_Morphometry_Protocols.pdf, at the time of writing).

## Data processing

### FreeSurfer

Cortical reconstruction and volumetric segmentation for all 134 images were performed with the FreeSurfer software suite, version 6.0.0 (http://surfer.nmr.mgh.harvard.edu/). We selected the T1 image with the least motion artefact (in case there were several attempts due to visible motion during scan) and then applied the “recon-all” processing stream with default parameters. It begins with transformation to Talaraich space, intensity inhomogeneity correction, bias field correction (Sled et al., 1998), and skull-stripping (Ségonne et al., 2004). Thereafter, WM is separated from GM and other tissues and the volume within the created WM–GM boundary is filled. After this, the surface is tessellated and smoothed. After these preprocessing steps are completed, the surface is inflated (Fischl, Sereno, & Dale, 1999) and registered to a spherical atlas. This method adapts to the folding pattern of each individual brain, utilizing consistent folding patterns such as the central sulcus and the sylvian fissure as landmarks, allowing for high localization accuracy (Fischl, Sereno, Tootell, et al., 1999). FreeSurfer uses probabilistic approach based on Markov random fields for automated labeling of brain regions. Cortical thickness is calculated as the average distance between the WM–GM boundary and the pial surface on the tessellated surface (Fischl & Dale, 2000). The cortical thickness measurement technique has been validated against post-mortem histological (Rosas et al., 2002) and manual measurements (Kuperberg et al., 2003; Salat, 2004).

### FreeSurfer manual edits and the FinnBrain quality control protocol

We used Freeview to view and edit the images using the standard command recommended by the FreeSurfer instructions with the addition of the Desikan–Killiany atlas that allowed us to correctly identify the ROIs where errors were found. Images with excess motion artefact or large unsegmented regions (extending over multiple gyri, examples provided in Supplementary Figure 1) were excluded. The images that passed the initial quality check were then manually edited and after that we ran the automated segmentation process again as suggested by FreeSurfer instructions. The images were then inspected again for errors, and the ROIs with errors were excluded in the FinnBrain quality control protocol, either from CT analyses (in cases where errors affected WM–GM or pial borders) or from volumetric analyses (all ROIs that were excluded from CT analyses, and ROIs with errors in cortical labeling).

#### Errors in borders

The automatically segmented images generated by FreeSurfer software suite were visually inspected and the found errors were either manually corrected or the ROI with the error was simply excluded depending on the type of error. Excess skull fragments were removed where the pial border was affected by them (Figures 1a and 1b). Arteries were removed to avoid segmentation errors between arteries and WM (especially relevant for anterior temporal areas and the insulae). This was done by setting the eraser to only delete voxels with intensity between 130 and 190 in the brainmask volume. The arteries were removed throughout the image with no regard to whether they caused issues in the segmentation on that specific slice. An example can be seen in Figure 1c.

**Figure 1.**
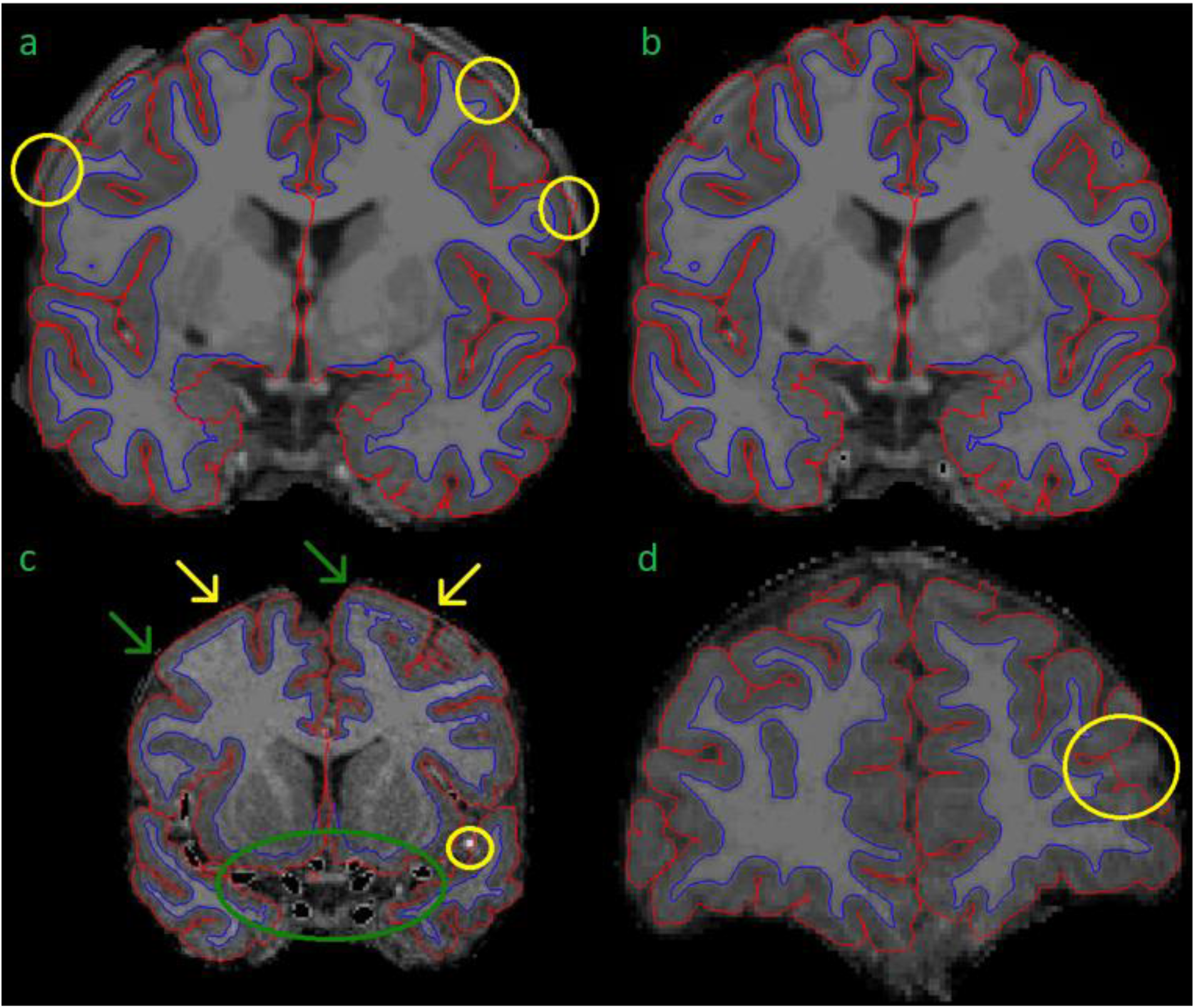
Figure 1 presents some common errors and fixes related to the pial border and non-brain tissues. Figure 1a demonstrates how skull fragments can cause errors in pial border (yellow circles). Figure 1b presents the same subject with skull fragments removed. In Figure 1c, arteries were removed (green circle). We removed voxels with an intensity between 130 and 190, and therefore some parts of arteries were not removed (yellow circle). Figure 1c also demonstrates the challenges with artefact, meninges, and the pial border. In some areas, the pial border may extend into the meninges (yellow arrows). Meanwhile, at the other end of the same gyrus, the border may seem correct (green arrows). It is difficult to fix these errors manually. Additionally, the visible motion artefact adds further challenges to manual edits of the pial border. In Figure 1d, the pial border cuts through a gyrus.

One typical error was that parts of the superior sagittal sinus (SSS) were included within the pial border. We stopped editing the SSS after an interim assessment as it was an arduous task with little effect on final results. All information regarding SSS edits is presented in Supplementary materials SSS.

In addition, there were errors that could not be fixed easily. In some cases, the pial border may cut through the cortex (Figure 1d shows an error in the left rostral middle frontal region). In these cases, the remaining GM mask is too small, and this error cannot be easily fixed in Freeview. Manual segmentation of a T1 image is labor intensive and hard to conduct reliably with 1 mm^3^ resolution even when the edits would cover small areas. Moreover, the FreeSurfer instructions do not recommend this approach. Additionally, the WM mask edits recommended in FreeSurfer instructions would not fix all cases where the cortical segmentation is too thin, as the WM mask often seemed adequate in these areas (an example presented in Supplementary Figure 2). Therefore, we simply had to exclude the ROI(s) in question.

Small errors of the WM–GM border were prevalent throughout the brain. To find and correct these, we systematically examined the brain from all three directions one hemisphere at the time. The corrections were made by erasing excess WM mask. This process is demonstrated in Figure 2. WM–GM border was inspected after the manual edits. A continuous error of at least ten slices in the coronal view led to exclusion of all the ROIs directly impacted by the error. Furthermore, ubiquitous errors in the WM–GM border, as markers of motion artefact, led to exclusion of the whole brain (as in Figure 3).

**Figure 2.**
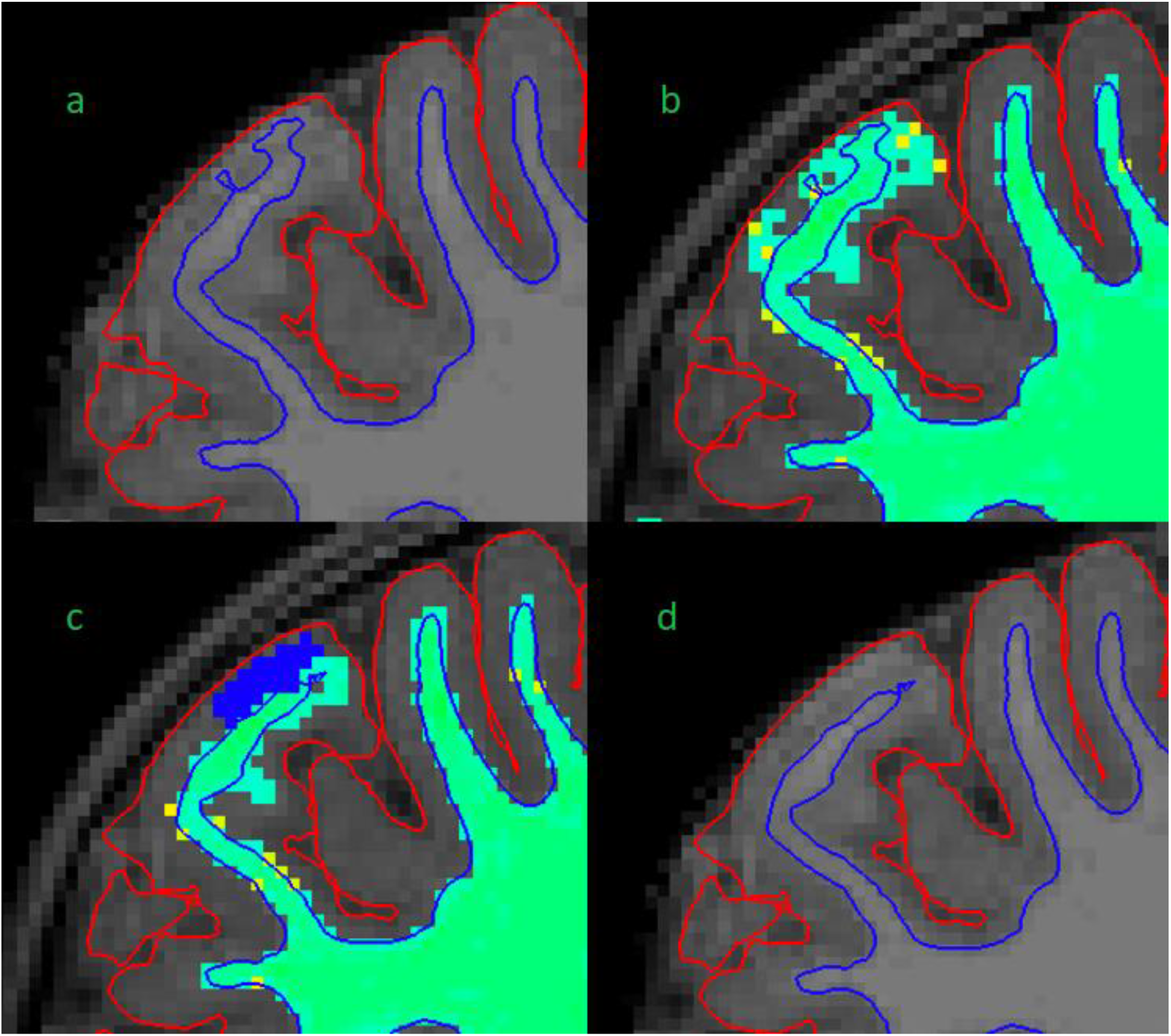
Figure 2 demonstrates our white matter (WM) mask editing protocol. Figure 2a shows a typical error in the border between white and gray matter (WM–GM border), where it extends too close to the pial border. Errors such as this are searched for in the “brainmask” volume (Figures 2a and 2d). Figure 2b shows the same error in “wm” volume with “Jet” colormap (Figures 2b and 2c). Figure 2c shows how we fixed these errors by erasing the erroneous WM mask (blue voxels). Figure 2d shows the final result after the second recon-all.

**Figure 3.**
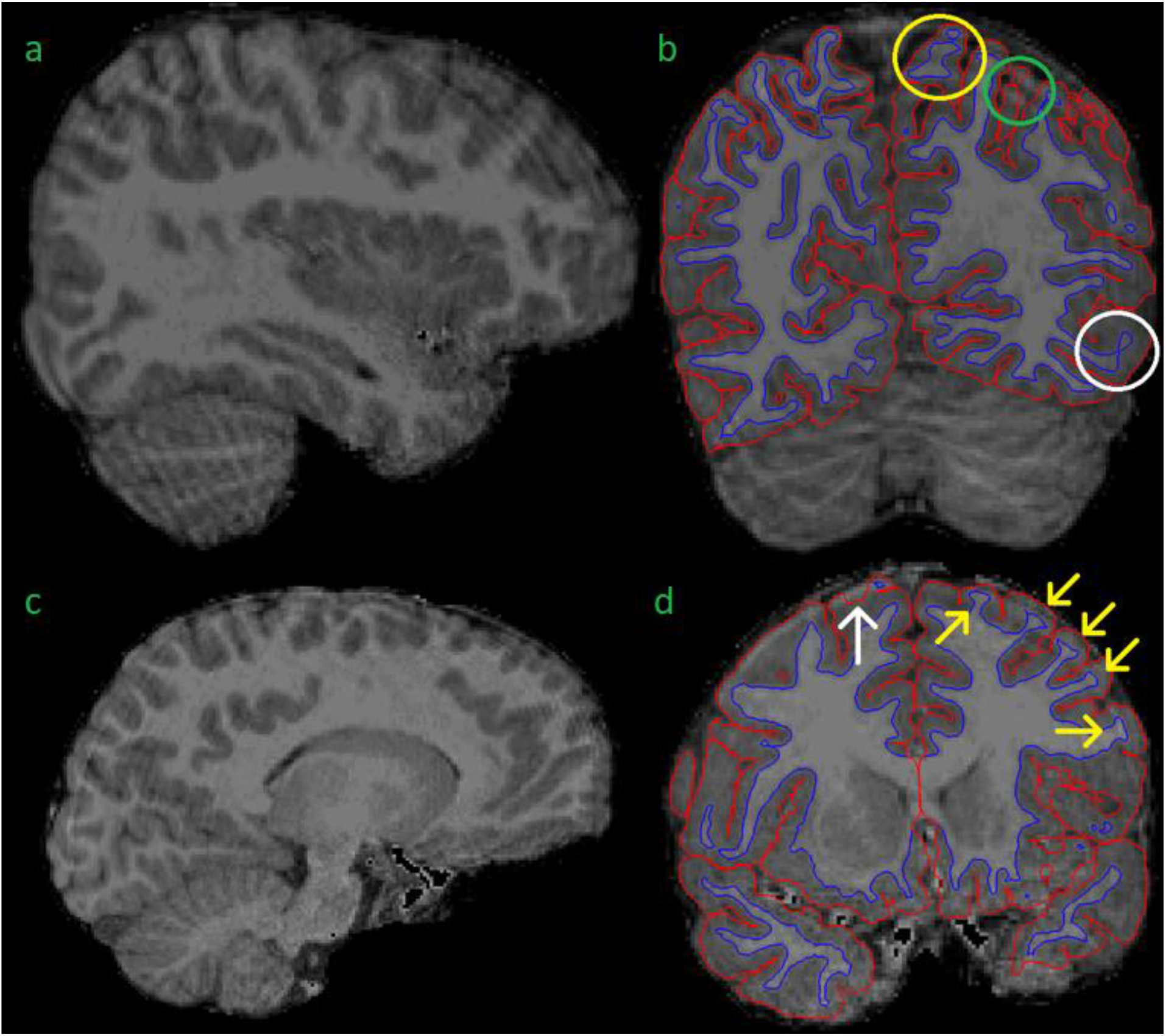
Figure 3 presents some examples of excluded brain images. Figure 3a shows “waves” throughout the image, marking motion artefact. Figure 3b shows the same subject as in 3a in a coronal view and borders visible. This image shows motion artefact related errors in the border between white and gray matter (WM–GM border), denoted by the yellow circle. Additionally, there is potential unsegmented area due to motion artefact (green circle) and poor contrast between WM and GM (white circle). Figures 3c and 3d show another excluded subject. The motion artefact in 3c is not as pronounced as in 3a. However, 3d still shows some typical errors for images with much artefact. There is a clear pial error (white arrow). Additionally, the yellow arrows show typical cases, where the “ringing” causes the WM mask to “widen” where the actual WM meets the ringing motion artefact.

Furthermore, there are some error types that cannot be easily fixed but also do not warrant exclusion. One such problem is that the pial border often extends into the cerebrospinal fluid or meninges around the brain (Supplementary Figure 3). The issue with this type of error is that sometimes the real border between GM and the surrounding meninges cannot be denoted visually and therefore the error cannot be reliably fixed. This problem is further complicated by the fact that motion artefact may mimic the border between GM and meninges making the visual quality control challenging (Figure 1c and Supplementary Figure 4).

There were some minor incongruities in multiple images. A common example can be seen in Supplementary Figure 5, where there seems to be a potential error in the pial border. Areas like this look normal in other planes. A less common example is shown in Supplementary Figure 6, where there is an apparent discontinuation in the WM–GM border. Similarly, there was no discontinuation in other planes. Both these minor incongruities were considered partial volume effects related to the presentation of a 3D surface in 2D slices. Therefore, both cases were included.

#### Errors in cortical labeling

A common issue was the presence of WM hypointensities in the segmented images. They sometimes appeared in the cortex, and they led to exclusion from volumetric analyses in the FinnBrain quality control protocol if there was a continuous error of at least two voxels in at least two directions (non-diagonally). These errors were typically small and did not cause errors in pial or WM–GM borders (Supplementary Figure 7). The hypointensities themselves were rarely successfully fixed by editing the WM mask and therefore were left unedited unless they caused errors in the GM–WM border. We tried to fix the errors in the WM–GM border and when unsuccessful, we simply excluded the ROI in question from both CT and volumetric analyses (Figures 4a and 4b). Of note, these errors can only be seen with the anatomical labels as overlays, unless they affect the WM–GM border.

**Figure 4.**
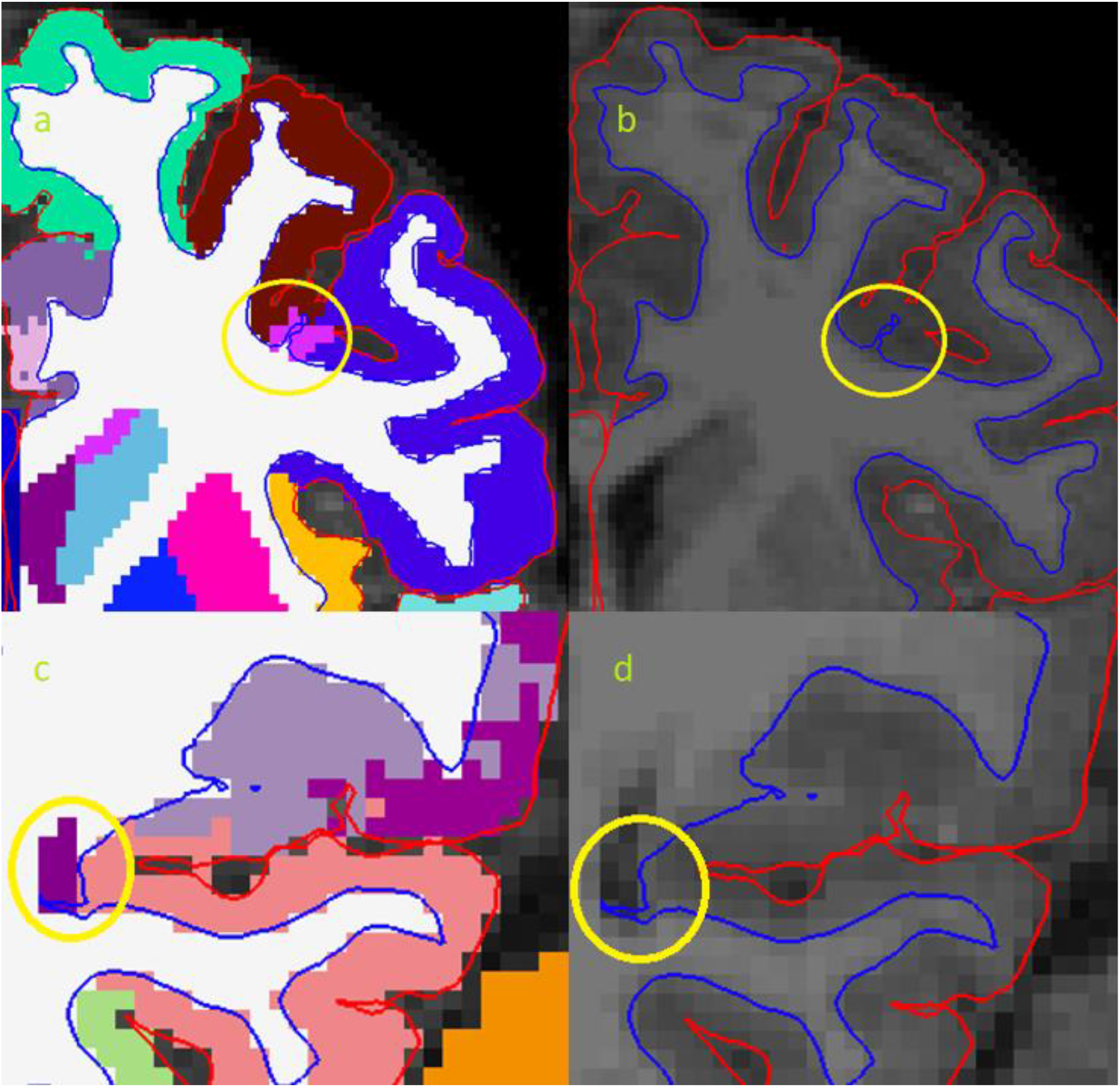
Figures 4a and 4b show a white matter (WM) hypointensity that affects the border between white and gray matter (WM–GM border), denoted by a yellow circle. Figures 4c and 4d show how the posterior part of the lateral ventricle causes distortion to the WM–GM border (yellow circle).

One typical error occurred at the posterior end of the lateral ventricles, where it may cause segmentation errors in the adjacent cortical regions, typically the precuneus and the lingual gyrus. These regions were excluded from volumetric analyses in the FinnBrain quality control protocol when the lateral ventricle or WM hypointensities appeared between the GM–WM border and the pial border (at least two adjacent voxels in at least two directions [non-diagonally], the voxels in question are not required to be fully between the borders) (Supplementary Figure 8), and from CT analyses when there was a distortion in the GM–WM border (Figures 4c and 4d). Unfortunately, hypointensities often appeared in ROI junctions, leading to exclusion of multiple regions due to one error (Supplementary Figure 9). Similar errors were seen in the ENIGMA protocol as well (Supplementary Figure 10).

#### Errors in subcortical labeling

Putamen was often mislabeled by FreeSurfer in our sample. Errors were addressed by adding control points, but the edits were largely unsuccessful. Consequently, we are currently working on separately validating subcortical segmentation procedures for our data. All information regarding the subcortical labeling is presented in Supplementary materials Subcortex.

### ENIGMA quality control protocol

After the quality control that entailed manual edits, we conducted a quality check with the ENIGMA Cortical Quality Control Protocol 2.0 (April 2017) (http://enigma.ini.usc.edu). Therein, the FreeSurfer cortical surface measures were extracted and screened for statistical outliers using R (https://www.r-project.org/) and visualized via Matlab (Mathworks) and bash scripts. Visual representations of the external 3D surface and internal 2D slices were generated and visually inspected according to the instructions provided by ENIGMA in https://drive.google.com/file/d/0Bw8Acd03pdRSU1pNR05kdEVWeXM/view (at the time of writing). The ENIGMA Cortical quality check instructions remark how certain areas have a lot of anatomical variation and therefore they note the possibility to be more or less stringent in their quality control. Considering this and the fact that the example images provided in the ENIGMA instructions are limited in number and as such cannot show every variation, we deemed necessary to describe how we implemented these instructions in our sample.

#### The external view

We started by viewing the external image. The pre- and postcentral gyri were assessed for meninge overestimations, which can manifest as “spikes” (Supplementary Figure 11a) or flat areas (Supplementary Figure 11b). These error types were rare in our sample. These cases were excluded as instructed.

The supramarginal gyrus has a lot of anatomical variability and when quality checking it, we decided to be lenient as suggested by the ENIGMA instructions. We only excluded cases where the border between supramarginal and inferior parietal regions cuts through a gyrus, leading to discontinuous segments in one of the regions (Figure 5a). In some rare cases, this type of error also happened with the postcentral gyrus (Supplementary Figure 12), and these cases were also excluded. Similarly, in cases with supramarginal gyrus overestimation into the superior temporal gyrus, we only excluded clear errors (examples presented in Supplementary Figure 13).

**Figure 5.**
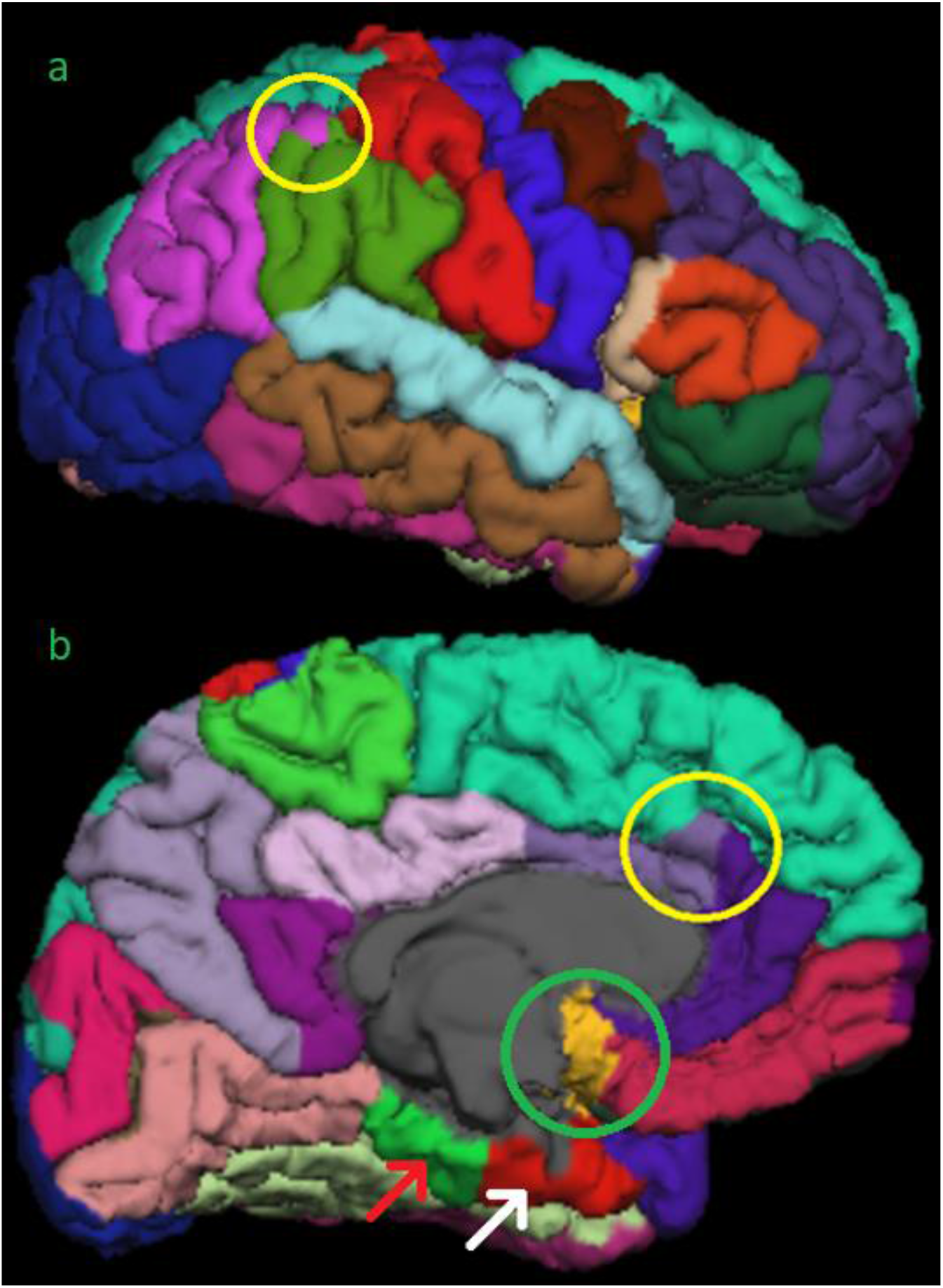
Figure 5a shows an error (yellow circle) where the inferior parietal area (purple) cuts through a whole gyrus in the supramarginal region (green). This area has a lot of variation and only clear errors led to exclusion in our ENIGMA quality control protocol. Figure 5b shows insula overestimation in the midline (green circle). Furthermore, the poor image quality can be seen the areas adjacent to the base of the skull, such as parahippocampal (green area denoted by a red arrow) and entorhinal (red area denoted be a white arrow). Additionally, there is an error in the border between superior frontal and caudal anterior cingulate. This border should follow the sulcal line. The rostral anterior cingulate was not considered erroneous in these cases.

One commonly seen error is insula overestimation into the midline (Figure 5b). In these cases, we exclude insula and the region(s) adjacent to it in the midline (e.g., the medial orbitofrontal region in the case of Figure 5b).

The border between the superior frontal region and the cingulate cortex (Figure 5b and Supplementary Figure 14) is one typical place for errors. A prominent paracingulate sulcus, that is more common on the left than on the right hemisphere, may cause underestimation of the cingulate cortex and consequently overestimation of the superior frontal region. This was typically seen on the left caudal anterior cingulate (Figure 5b), where we excluded the cases where the border did not follow sulcal lines anteriorly (as was demonstrated in the image examples in the instructions). In rare cases the border between posterior cingulate and superior frontal region was affected (Supplementary Figure 14), and these were also excluded. The pericalcarine region was overestimated in some cases. According to the instructions cases where the segmentation is confined to the calcarine sulcus should be accepted. Therefore, we excluded cases where the pericalcarine region extended over a whole gyrus into the lingual gyrus or the cuneus. An example can be seen in Supplementary Figure 15.

Cases of superior parietal overestimation were excluded as instructed. These errors were rare in our sample. Similarly, errors in the banks of the superior temporal sulcus were excluded as instructed.

The border between the middle and inferior temporal gyrus was not assessed, as the instructions suggested that most irregularities seen there are normal variants or relate to the viewing angle.

Similarly, we did not quality check the entorhinal/parahippocampal regions in the external view, as there is a lot of variation in the area. The ENIGMA instructions describe underestimations in 70–80% of cases. Furthermore, this region looks poor in practically all images (e.g., in Figure 5b) as do all the regions adjacent to the base of the skull and therefore, in our opinion, the quality assessment in those regions requires additional procedures, that are beyond the scope of the current study, to confirm their usability in statistical analyses.

#### The internal view

In the internal view, regions with unsegmented GM were excluded. These errors often reflect WM hypointensities seen in Freeview (Supplementary Figure 10). Interestingly, even quite large hypointensities do not necessarily equate to errors in the borders set by FreeSurfer and therefore do not always have an adverse effect on CT calculations.

Temporal pole underestimations were sometimes seen. However, the cases were rarely as clear as presented in the instructions. Therefore, we had to use both coronal and axial views to assess the situation and make exclusions when both views supported an error in segmentation.

One of the errors commonly seen in our sample was the erroneous segmentation in the lateral parts of the brain. This was particularly prevalent in the middle temporal gyri (Supplementary Table 2). This error was assessed from 2D slices, wherein what seems to be an error may be caused by partial volume effects. For example, in Supplementary Figure 16a, there seems to be a possible error on the right middle temporal region. If we look at the same image in Freeview, the same position seems to be segmented normally, especially when confirmed in the axial view (Supplementary Figures 16b and 16c). Consequently, we only made exclusion when clear errors were seen in two adjacent slices. Particularly clear example of this can be seen in Figure 6, where the WM extends outside the segmentation. The error is also visible in the external view, where these regions do not appear as smooth as normally (Supplementary Figure 17), however the decisions to exclude a ROI were always made based on the internal view. This kind of error was significantly harder to recognize in Freeview and represents the most striking difference in results between the ENIGMA and FinnBrain quality control protocols.

**Figure 6.**
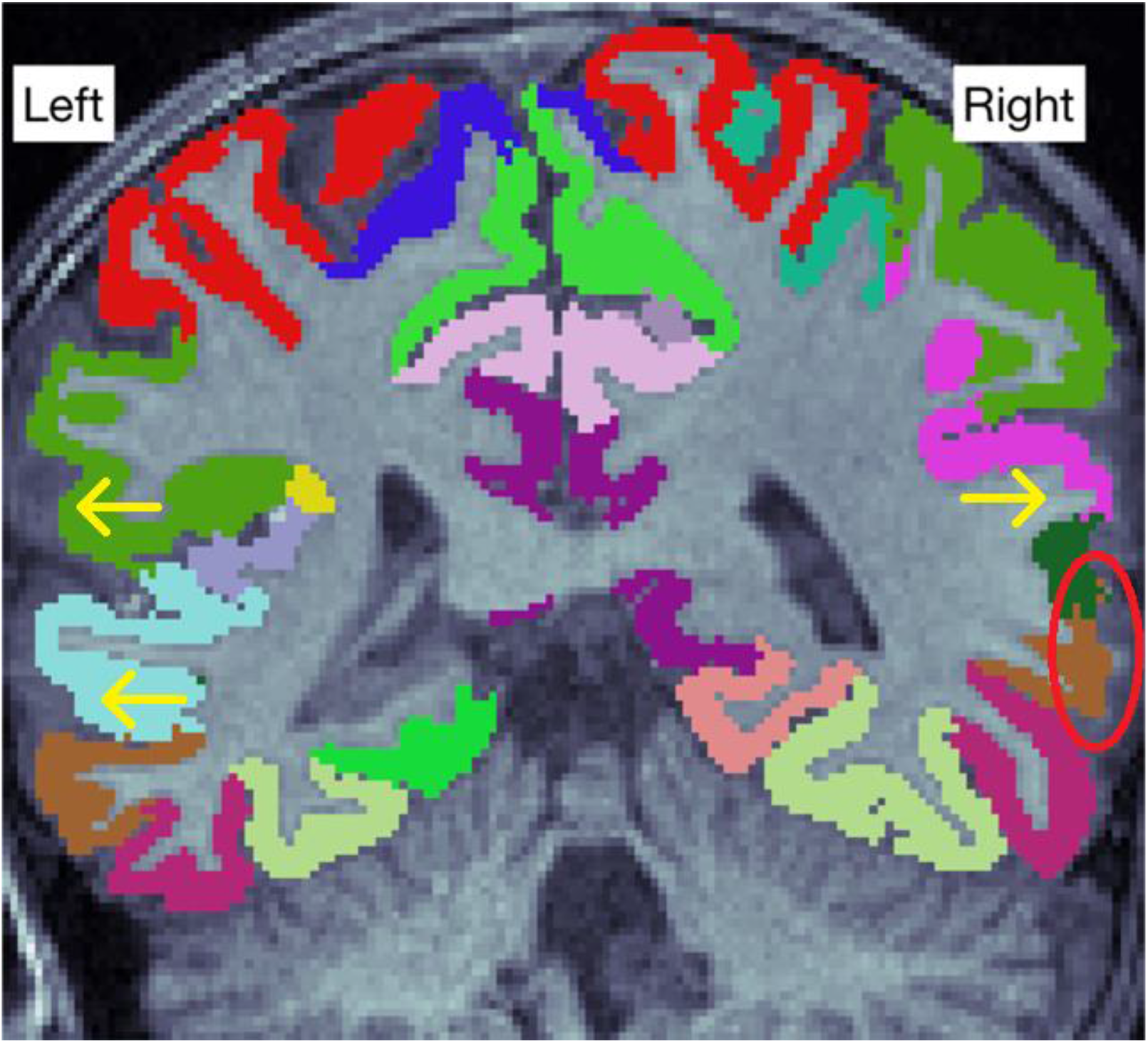
Figure 6 shows errors in the lateral parts of the image (arrows). An especially clear error is denoted by the red circle, where some white matter is seen outside the cortical segmentation.

#### Statistical outliers

After the systematic viewing of all the problem regions, we inspected the statistical outliers. This rarely led to new exclusions, as many of the statistical outliers were among the excluded subjects or the outliers were ROIs where the instructions did not give any tools to assess whether they were correct. Therefore, we had to simply double check the internal view to rule out segmentation errors.

### Exclusions

We decided to use a dichotomous rating scale: pass or fail. The amount of motion artefact (marked by “concentric rings” or “waves”) and the clarity of the WM–GM border were assessed from the original T1 image. In borderline cases, we ran the standard recon-all and made new assessment based on the segmented image. Massive segmentation errors such as large missing areas or ubiquitous errors in WM–GM border were reasons for exclusion. Additionally, ENIGMA exclusion criteria were implemented as instructed. In some borderline cases, another expert rater assessed the image quality and agreement was reached to either include or exclude the image. Some images that were considered for inclusion but excluded after the first recon-all can be seen in Figure 3. These images had significantly more artefact than other images in our sample, although arguably they could have been included since the amount of artefact could be described as “moderate”. However, we decided to implement strict exclusion criteria to ensure high quality of data.

### Alternate processing: Optional registration flags in FreeSurfer

We compared the FreeSurfer default recon all to recon-all with the “-mprage” and “-schwartzya3t-atlas” optional flags. All information regarding optional flags analyses is presented in Supplementary materials Optional flags.

### Alternate processing: CAT12

A previous study conducted in the elderly demonstrated good agreement between FreeSurfer and CAT12 estimates of CT (R^2^= 0.83), although CAT12 produced systematically higher values than FreeSurfer (Seiger et al., 2018). Therefore, we decided to explore the agreement between the two software in a pediatric population. All information regarding CAT12 analyses is presented in Supplementary materials CAT12.

### Statistics

Statistical analyses were conducted using the IBM SPSS Statistics for Windows, version 25.0 (IBM Corp., Armonk, NY, USA). The ROI data was confirmed to be normally distributed using JMP Pro 15 (SAS Institute Inc., Cary, NC) based on visual assessment and the similarity of mean and median values.

To compare the differences between the included and excluded groups, we performed independent samples t-tests for age from birth at scan, gestational age at scan, gestational age at birth, birthweight, maternal age at term, and maternal body mass index (BMI) before pregnancy. In addition, we conducted Chi-Square tests for child gender, maternal education level (three classes: 1 = Upper secondary school or vocational school or lower, 2 = University of applied sciences, 3 = University), maternal monthly income estimate after taxes (in euros, four classes: 1 = 1 500 or less, 2 = 1 501 to 2 500, 3 = 2 501 to 3 500, 4 = 3 501 or more), maternal alcohol use during pregnancy (1 = yes, continued to some degree after learning about the pregnancy, 2 = yes, stopped after learning about the pregnancy, 3 = no), maternal tobacco smoking during pregnancy (1 = yes, continued to some degree after learning about the pregnancy, 2 = yes, stopped after learning about the pregnancy, 3 = no), maternal history of disease (allergies, depression, asthma, anxiety disorder, eating disorder, chronic urinary tract infection, autoimmune disorder, hypercholesterolemia, and hypertension), maternal medication use at gestational week 14 (non-steroidal anti-inflammatory drugs, thyroxin, selective serotonin reuptake inhibitor [SSRI] or serotonin–norepinephrine reuptake inhibitor [SNRI], and corticosteroids), or at gestational week 34 (thyroxin, SSRI or SNRI, corticosteroids, and blood pressure medications). The categories in history of disease and medication during pregnancy were only included in statistical analyses, when there were at least four participants that had history of the disease or used the medication (to limit the chance of false positives).

The inclusion criterion for the ROI based comparisons was passing the ENIGMA quality control protocol. To compare edited FreeSurfer to unedited FreeSurfer, we conducted a paired samples t-test. We calculated the absolute values of the change in CT between unedited and edited images for each ROI separately using the following formula: (C_D_/C_U_) * 100%, where C_D_ is the absolute value of the difference in mean CT between edited and unedited images and C_u_ is the mean CT in the unedited images. Furthermore, we conducted a paired samples t-test with the mean CT values from all ROIs to measure the change between edited and unedited images.

All significances were calculated 2-tailed (α = 0.05). To adjust for multiple comparisons in ROI-based analyses, we conducted the Bonferroni correction by setting the p value to 0.05 divided by the number of comparisons (= the number of ROIs = 68), resulting in p = 0.000735. We notify that the p value cut off for the current study is somewhat arbitrary and thus we also report the raw p values in the tables.

## Results

### Demographics

There were no significant differences between the included and excluded subjects’ age from birth at scan, gestational age at scan, gestational age at birth, birth weight, maternal age at term, maternal education level, maternal monthly income, maternal history of disease, maternal alcohol use during pregnancy, or maternal tobacco smoking during pregnancy. There was a significant difference in maternal BMI before pregnancy (p = 0.03). In the included group, mean maternal BMI was 23.9 (n = 121) vs. 26.0 in the excluded group (n = 24, information from one participant missing). Two types of medication were more common in the excluded group: SSRI or SNRI medication at 14 gestational weeks (p = 0.03; included group 109 no, 3 yes; excluded group 20 no, 3 yes) and blood pressure medication at 34 gestational weeks (p = 0.03; included group 113 no, 3 yes; excluded group 21 no, 3 yes). In addition, there was a marginally significant difference in SSRI/SNRI use at 34 gestational weeks (p = 0.06; included group 112 no, 4 yes; excluded group 21 no, 3 yes). Of note, these results are not optimal to determine whether the listed early exposures are associated with poorer image quality as such but such comparisons may be useful to conduct before final analyses in any data set (and are also included for descriptive purposes)(please see related articles: Buss et al., 2012; Chen et al., 2014; Edlow, 2017; Morales et al., 2018; A. Rodriguez et al., 2008; Alina Rodriguez, 2010; Tanda & Salsberry, 2014).

### Comparison between unedited and manually edited FreeSurfer segmentations

The difference in CT was not significant after Bonferroni correction in 57/68 (83.8%) regions. Unedited images had significantly larger CT values in 2/68 (2.9%) regions: the right rostral anterior cingulate and right superior temporal regions. Edited images had significantly larger CT values in 9/68 (13.2%) regions: the left and right caudal middle frontal, left and right inferior temporal, left and right superior parietal, right precentral, right superior frontal, and right supramarginal regions. The smallest (both absolute and relative) change was observed in the left rostral middle frontal (0.0003mm, 0.011%) and the largest (both absolute and relative) in the right caudal middle frontal (0.0526mm, 1.857%) region. The CT changes and raw p-values for all ROIs are presented in Supplementary Table 3.

The mean change in absolute CT values between the unedited and edited images was 0.0129mm (0.441%). When we include the direction of the change in the analysis, edited images had higher CT values (mean 0.00264mm, 0.0901%), although the difference was not statistically significant (p = 0.217).

Pearson correlations between edited and unedited images were calculated by ROI, they all were positive and ranged from 0.725 in the left insula to 0.984 in the left banks of the superior temporal sulcus region. All remained statistically significant after Bonferroni correction. The correlations are displayed in Supplementary Table 4.

### The ENIGMA and FinnBrain quality control protocols

Overall, the FinnBrain CT quality check protocol was the most permissive with 7824 accepted datapoints out of possible 8228 (FinnBrain volume 7341, ENIGMA 7208). The largest differences in both directions between FinnBrain CT and ENIGMA quality check protocols were found in left middle temporal (FinnBrain CT 119; ENIGMA 77; difference 42) and left precuneus (FinnBrain CT 91; ENIGMA 110; difference 19). The worst quality areas (measured by total datapoints across all 3 protocols) were the left precuneus and the left lingual gyrus, with 260 and 263 datapoints, respectively. The number of subjects with an error in a specific ROI can be seen in Supplementary Table 2. The number of data sets that passed the protocols with no ROI exclusions was relatively low: three for the FinnBrain volumetric protocol, 22 for the FinnBrain CT protocol, and three for the ENIGMA protocol (15 passes for the external and 25 passes for the internal view; notably, the internal was rated as “pass” if it did not result in additional exclusions when viewed after the external view, and therefore the number of passes is overestimated).

## Discussion

In this article, we described the semiautomated segmentation procedure we used for image processing in detail. While this work relied heavily on existing guidelines by FreeSurfer and the ENIGMA consortium, we believe this article is of help for investigators that are new to pediatric neuroimaging. We add to the existing literature by assessing the effects of our manual edits on CT values, reporting the agreement between FreeSurfer and CAT12, and comparing the FreeSurfer’s standard recon-all to other optional flags.

The manual edits had minor effects on the CT values, less than 2% in all regions (comparable with earlier results by McCarthy et al., 2015), and the differences were rarely significant. Edited images had larger CT values in most cases where significant differences were seen. This is not surprising, as most of the editing time is spent correcting small errors in the WM–GM border, and fixing these errors typically thickens the cortex (as can be seen in Figure 2). The opposite effect was seen in only two regions. In case of the right rostral anterior cingulate, there are some arteries adjacent to it, and erasing them may have had a thinning effect on cortical thickness (Supplementary Figure 18). However, the reason for the apparent thinning of the superior temporal region is unclear.

For inclusion and exclusion criteria of images, we propose that there are two major approaches: micro and macro scale approach. In the micro scale assessment, we could find the errors as described in the methods section and score the image based on their number and size. However, this approach has multiple challenges. Firstly, there are many errors that do not warrant exclusion of the ROI in question, e.g., small errors in the WM–GM border (demonstrated in Figure 2a). In some cases, these types of errors were abundant despite rarely meeting the exclusion criteria. How should the number of these errors be calculated and what weight should they be given compared to larger errors? Secondly, in many cases it is not obvious whether there is an error in the slice or not (one typical case is an image with poor WM–GM contrast). If we were to count errors by the number of slices with a certain type of error, differences between raters could lead to large differences in these cases. These could be viewed by multiple expert raters and discussed, however that would be very time consuming and arduous, while one of the main goals of semiautomated segmentation programs is to make the process as fast and easy as possible. Thirdly, quality control protocols are often described on a general level in scientific studies (Barnes-Davis et al., 2020; Boutzoukas et al., 2020; El Marroun et al., 2016; Kamson et al., 2016), and therefore there is no commonly accepted way to assess all the errors in the automated segmentation.

In contrast, in the macro scale assessment, the rater can quickly look at the brain image, and assess the amount of motion artefact (marked by “waves” or “concentric rings” in the image) and the clarity of the WM–GM border. In borderline cases, the image can be segmented and then assessed for major segmentation errors, such as ubiquitous errors in the WM–GM border or large unsegmented areas. One key challenge with this approach is the lack of objective criteria, as these types of errors are very difficult to quantify or to describe in articles or instructions. However, the expert rater makes this same assessment for all images and can learn to quickly exclude the images that are of significantly poorer quality than others, and therefore a high internal reliability should be attainable. Furthermore, as this type of assessment can be made quickly, unclear cases can be verified by other raters with little additional time commitment. Considering the pros and cons of both approaches, we decided to use macro scale assessment for exclusion of whole images. Furthermore, we decided to apply it on a dichotomous pass or fail scale and skip further quality classification. One possible downside is the loss of subcategories in the accepted sample, since image quality can be included in regression analyses (Shaw et al., 2007). However, in our study, we perform a rigorous quality control protocol that rates image quality on a level of single ROIs, and therefore all datapoints in the final sample are of high quality. Consequently, we believe a further categorization based on overall image quality would not add significant value in this case.

We decided to apply the widely used and accepted ENIGMA quality control protocol (Thompson et al., 2020) to support decisions on inclusion and exclusion of ROIs. It has previously been implemented for both adults (Thompson et al., 2020) and children (Boedhoe et al., 2018; Hoogman et al., 2019). The internal view of ENIGMA protocol gives 16 slices with color coded segmented ROIs. This gives a good overall view of the brain, but it does not allow for exploration of unclear cases, and some errors can be completely missed if they are not located in the slices presented by ENIGMA. To explore this issue, we presented our own FinnBrain quality control protocol, and as a result of using Freeview for slice-by-slice assessment of the brain (e.g., the errors seen in Figure 4) it was more sensitive to certain types of small errors than the ENIGMA protocol. However, this protocol was not implemented for the final analyses due to the challenges discussed earlier in this article, such as the large number of minor errors and the lack of consensus on how to treat them. For example, the areas that were the most commonly excluded from the volumetric analyses in the FinnBrain protocol were the left lingual gyrus and left precuneus (Supplementary Table 2). Both regions are adjacent to the posterior tip of the lateral ventricle and were therefore often excluded due to a few mislabeled voxels. Overall, the FinnBrain protocol was more permissive than the ENIGMA protocol. One major reason for this is that it lacks the external view that ENIGMA has and therefore cannot assess errors in borders between ROIs. Therefore, even if the FinnBrain quality control protocol were implemented, it would have to be implemented together with the ENIGMA protocol. However, future studies should explore the utility of slice-by-slice assessment of the Freeview image, as some of the errors found via that method may be large enough to warrant exclusion from statistical analyses.

The key practical benefit in our manual edits protocol is the relative ease of application. Errors caused by skull fragments are very clear and easy to fix (Figure 1a). Fixing arteries by erasing all voxels between certain intensity values requires practically no decision-making during execution. Edits in the WM–GM border take the most time and require the most expertise. However, as the edits are followed by another automated recon-all protocol, that considers the human input in calculations, the editor cannot decide the exact delineation between WM and GM, and therefore cannot make errors that would mandate editing the image again from scratch. Such errors were possible while editing the SSS (please see Supplementary materials SSS), however SSS edits were stopped after an interim analysis. While it could be argued that the effect on CT values is not worth the time that manual edits require, we believe that systematic manual edits protocol has an additional benefit: It maximizes the chance to find and fix errors that would lead to exclusion of the ROI in question, therefore increasing the number of valid datapoints in the final sample.

There is an increased need for manual edits and diligent quality control in pediatric imaging. Children move more than adults during scans (Blumenthal et al., 2002; Poldrack et al., 2002; Theys et al., 2014), and therefore there is an increase in ringing and blurring artefacts in images. The artefact can lead to unreliable cortical parcellations, and the errors must be noted and fixed when possible. Furthermore, the choice of automated segmentation tool can be very influential. For example, in adults, FreeSurfer and CAT12 have shown good agreement (Masouleh et al., 2020; Seiger et al., 2018), however in our sample the agreement was relatively poor. CAT12 often overestimated CT compared to FreeSurfer, but the opposite was also true a significant number of cases, showing that the disagreement was not systematic. Therefore, the results cannot be reliably compared in this population. Please see Supplementary materials CAT12 for more discussion. Furthermore, a child’s brain undergoes non-linear regional developments through its development, which means it cannot simply be considered a slightly smaller adult brain (Phan, Smeets, et al., 2018; Wilke et al., 2003). Therefore, adult templates may be suboptimal for developing brains (Muzik et al., 2000; Phan, Sima, et al., 2018; Yoon et al., 2009) and custom pediatric templates should be considered.

One possible way to increase the feasibility of anatomic segmentation in children is the creation and use of custom templates. Multiple pediatric atlases have been created (Phan, Smeets, et al., 2018), some including 5-year-olds (Fonov et al., 2011; Wilke et al., 2003). Furthermore, it is possible to create templates using software packages such as the Template-O-Matic (Wilke et al., 2008), and those provided by ANTs (Avants et al., 2011) or IDEAgroup (https://www.med.unc.edu/bric/ideagroup/software/). Using pediatric atlases could produce more accurate results compared to the adult ones, and this may even be the case when the atlas is external (images from children of similar age from outside the study) as compared to internal (images from within the study) (Yoon et al., 2009). Certain typical errors can appear in unedited pediatric FreeSurfer images when applying adult templates, as presented in a review by Phan et al. (Phan, Smeets, et al., 2018). The review presents the errors in pial border that were often seen in the temporal regions in our sample. On the other hand, cerebellum was mislabeled only once in our final sample (Supplementary Figure 19). Additionally, we did observe erroneous automatic segmentation in the subcortical regions, and we are preparing an article regarding the manual segmentation of these areas (Lidauer et al., in preparation). Similarly, pediatric atlases have their own challenges. One key challenge with age specific atlases is that the required specificity regarding the age range is unclear. Multiple age specific atlases, some of them freely available, have been created for neonates and infants (Kuklisova-Murgasova et al., 2011; Shi et al., 2011), and they have shown good agreement with the “gold standard” manual segmentation (Oishi et al., 2011; Serag et al., 2012). The age ranges in these atlases may be very specific, e.g. covering the preterm neonates aged between 29 and 44 gestational weeks (Kuklisova-Murgasova et al., 2011). In comparison, in older pediatric populations, the age ranges may be a few years or even more than ten years (Fonov et al., 2011; Wilke et al., 2003). In addition, pediatric atlases have the challenges that atlases have in general, such as the specificity of the group (e.g., a certain disease) and ROIs. Considering the multitude of different options in pediatric atlases, their use may complicate comparisons between studies. Therefore, we decided to use the standard adult atlas with appropriate quality control measures to counter the challenges this approach has. We were generally satisfied with the cortical segmentation results, but it remains an important venue to develop and validate implemented in mainstream software such as FreeSurfer (de Macedo Rodrigues et al., 2015; Zöllei et al., 2020).

One of the key limitations in our study is the reliance on visual assessment in the quality control. Considering the inherent arbitrariness of the visual assessment of motion artefact, there is interest in developing automated quality assessment algorithms (White et al., 2018). An automated, objective estimate of the severity of the motion might allow us to set universal standards on the different categories of motion severity. There are some challenging key questions that would need to be resolved before the creation of a system to correct for motion artefact: 1) how much different levels of motion affect different aspects of brain morphology (Blumenthal et al. provide estimates of the decrease in volume in a seemingly nonlinear manner, as the change from moderate to severe artefact causes a major drop in volumes compared to the other classifications); and 2) are the effects similar throughout the brain or are there significant regional differences. Considering these challenges, more research is needed before the effects of motion artefact can be accounted for automatically. Another approach is to lessen motion artefact by adding prospective motion correction (PMC) to the T1-weighted imaging sequence (Ai et al., 2021). The benefit is clearest in images with a lot of motion artefact, while the cost is poorer performance in some quality control measures such as signal to noise ratio compared to a MPRAGE sequence without PMC (Ai et al., 2021). While implementation of PMC could improve the quality of our data, it would not remove the need a quality control protocol such as the one we presented in this article, and therefore the existence of this alternative imaging sequence does not impact our main findings. Although we opted for a quality control protocol that performs visual quality control on a level of individual ROIs, investigators may additionally benefit from using custom software to detect potentially low quality data (Klapwijk et al., 2019).

## Conclusion

There is no one “gold standard” processing method for pediatric images, are thus there is methodological variation between different studies. Pediatric images are inherently more susceptible for segmentation errors than adult images due to increased motion during scans and the potential mismatch with adult templates and segmentation tools. This highlights the need for rigorous quality control to ensure high quality data. We believe that detailed method descriptions are crucial for maximal transparency that helps comparisons between studies.

In this article we have described in detail the semiautomated segmentation protocol used in the FinnBrain Neuroimaging Lab, including manual edits and the implementation of the ENIGMA quality control protocol. We decided to use the standard recon-all without optional registration flags, as they did not provide additional benefits. Furthermore, we observed a surprisingly poor agreement between FreeSurfer and CAT12 output. Our semiautomated segmentation protocol provides high quality pediatric neuroimaging data and could help investigators working with similar data sets.

## Acknowledgements

The authors declare no conflicts of interest.

EPP was supported by the Päivikki and Sakari Sohlberg Foundation. VK was supported by The Finnish Cultural Foundation (Lastenlinnan säätiö) (recruitment, collection of the MRI data and writing the manuscript). HM was supported by The Finnish Cultural Foundation. SN was supported by the State Grants for Clinical Research (ERVA) and Signe and Ane Gyllenberg Foundation. RK was supported by the Academy of Finland (308252), Signe & Ane Gyllenberg Foundation, the Hospital District of Southwest Finland (State research grant). LK was supported by the State Grants for Clinical Research (ERVA) and Brain and Behavior Research Foundation, YI Grant #1956. JJT was supported by the Hospital District of Southwest Finland (State research grant), Turku University Foundation, Emil Aaltonen Foundation, and Alfred Kordellin Foundation (data collection and data analysis) as well as and Sigrid Jusélius Foundation (interpretation of the data and writing the manuscript).

We thank our research nurse Susanne Sinisalo for her expertise in study management and performing the scans with the investigators and all participated FinnBrain Families.

## References

Ai, L., Craddock, R. C., Tottenham, N., Dyke, J. P., Lim, R., Colcombe, S., Milham, M., & Franco, A. R. (2021). Is it time to switch your T1W sequence? Assessing the impact of prospective motion correction on the reliability and quality of structural imaging. NeuroImage, 226, 117585. https://doi.org/10.1016/j.neuroimage.2020.117585

Al Harrach, M., Rousseau, F., Groeschel, S., Wang, X., Hertz-pannier, L., Chabrier, S., Bohi, A., Lefevre, J., & Dinomais, M. (2019). Alterations in Cortical Morphology after Neonatal Stroke: Compensation in the Contralesional Hemisphere? Developmental Neurobiology, 79(4), 303–316. https://doi.org/10.1002/dneu.22679

Alexander-Bloch, A., Clasen, L., Stockman, M., Ronan, L., Lalonde, F., Giedd, J., & Raznahan, A. (2016). Subtle in-scanner motion biases automated measurement of brain anatomy from in vivo MRI. Human Brain Mapping, 37(7), 2385–2397. https://doi.org/10.1002/hbm.23180

Avants, B. B., Tustison, N. J., Song, G., Cook, P. A., Klein, A., & Gee, J. C. (2011). A reproducible evaluation of ANTs similarity metric performance in brain image registration. NeuroImage, 54(3), 2033–2044. https://doi.org/10.1016/j.neuroimage.2010.09.025

Barnea-Goraly, N., Weinzimer, S. A., Ruedy, K. J., Mauras, N., Beck, R. W., Marzelli, M. J., Mazaika, P. K., Aye, T., White, N. H., Tsalikian, E., Fox, L., Kollman, C., Cheng, P., & Reiss, A. L. (2014). High success rates of sedation-free brain MRI scanning in young children using simple subject preparation protocols with and without a commercial mock scanner-the Diabetes Research in Children Network (DirecNet) experience. Pediatric Radiology, 44(2), 181–186. https://doi.org/10.1007/s00247-013-2798-7

Barnes-Davis, M. E., Williamson, B. J., Merhar, S. L., Holland, S. K., & Kadis, D. S. (2020). Extremely preterm children exhibit altered cortical thickness in language areas. Scientific Reports, 10(1), 1–10. https://doi.org/10.1038/s41598-020-67662-7

Beelen, C., Phan, T. V., Wouters, J., Ghesquière, P., & Vandermosten, M. (2020). Investigating the Added Value of FreeSurfer’s Manual Editing Procedure for the Study of the Reading Network in a Pediatric Population. Frontiers in Human Neuroscience, 14, 143. https://doi.org/10.3389/fnhum.2020.00143

Black, J. M., Tanaka, H., Stanley, L., Nagamine, M., Zakerani, N., Thurston, A., Kesler, S., Hulme, C., Lyytinen, H., Glover, G. H., Serrone, C., Raman, M. M., Reiss, A. L., & Hoeft, F. (2012). Maternal history of reading difficulty is associated with reduced language-related gray matter in beginning readers. NeuroImage, 59(3), 3021–3032. https://doi.org/10.1016/j.neuroimage.2011.10.024

Blumenthal, J. D., Zijdenbos, A., Molloy, E., & Giedd, J. N. (2002). Motion artifact in magnetic resonance imaging: Implications for automated analysis. NeuroImage, 16(1), 89–92. https://doi.org/10.1006/nimg.2002.1076

Boedhoe, P. S. W., Schmaal, L., Abe, Y., Alonso, P., Ameis, S. H., Anticevic, A., Arnold, P. D., Batistuzzo, M. C., Benedetti, F., Beucke, J. C., Bollettini, I., Bose, A., Brem, S., Calvo, A., Calvo, R., Cheng, Y., Cho, K. I. K., Ciullo, V., Dallaspezia, S.,… Van Den Heuvel, O. A. (2018). Cortical abnormalities associated with pediatric and adult obsessive-compulsive disorder: Findings from the enigma obsessive-compulsive disorder working group. American Journal of Psychiatry, 175(5), 453–462. https://doi.org/10.1176/appi.ajp.2017.17050485

Boutzoukas, E. M., Crutcher, J., Somoza, E., Sepeta, L. N., You, X., Gaillard, W. D., Wallace, G. L., & Berl, M. M. (2020). Cortical thickness in childhood left focal epilepsy: Thinning beyond the seizure focus. Epilepsy and Behavior, 102, 106825. https://doi.org/10.1016/j.yebeh.2019.106825

Buss, C., Entringer, S., Davis, E. P., Hobel, C. J., Swanson, J. M., Wadhwa, P. D., & Sandman, C. A. (2012). Impaired Executive Function Mediates the Association between Maternal Pre-Pregnancy Body Mass Index and Child ADHD Symptoms. PLoS ONE, 7(6), e37758. https://doi.org/10.1371/journal.pone.0037758

Chen, Q., Sjolander, A., Langstrom, N., Rodriguez, A., Serlachius, E., D’Onofrio, B. M., Lichtenstein, P., & Larsson, H. (2014). Maternal pre-pregnancy body mass index and offspring attention deficit hyperactivity disorder: a population-based cohort study using a sibling-comparison design. International Journal of Epidemiology, 43(1), 83–90. https://doi.org/10.1093/ije/dyt152

Clark, K. A., Helland, T., Specht, K., Narr, K. L., Manis, F. R., Toga, A. W., & Hugdahl, K. (2014). Neuroanatomical precursors of dyslexia identified from pre-reading through to age 11. Brain, 137(12), 3136–3141. https://doi.org/10.1093/brain/awu229

Dale, A. M., Fischl, B., & Sereno, M. I. (1999). Cortical surface-based analysis: I. Segmentation and surface reconstruction. NeuroImage, 9(2), 179–194. https://doi.org/10.1006/nimg.1998.0395

de Macedo Rodrigues, K., Ben-Avi, E., Sliva, D. D., Choe, M., Drottar, M., Wang, R., Fischl, B., Grant, P. E., & Zöllei, L. (2015). A FreeSurfer-compliant consistent manual segmentation of infant brains spanning the 0-2 year age range. Frontiers in Human Neuroscience, 9(FEB), 21. https://doi.org/10.3389/fnhum.2015.00021

Edlow, A. G. (2017). Maternal obesity and neurodevelopmental and psychiatric disorders in offspring. Prenatal Diagnosis, 37(1), 95–110. https://doi.org/10.1002/pd.4932

El Marroun, H., Tiemeier, H., Franken, I. H. A., Jaddoe, V. W. V., van der Lugt, A., Verhulst, F. C., Lahey, B. B., & White, T. (2016). Prenatal Cannabis and Tobacco Exposure in Relation to Brain Morphology: A Prospective Neuroimaging Study in Young Children. Biological Psychiatry, 79(12), 971–979. https://doi.org/10.1016/j.biopsych.2015.08.024

Epstein, J. N., Casey, B. J., Tonev, S. T., Davidson, M., Reiss, A. L., Garrett, A., Hinshaw, S. P., Greenhill, L. L., Vitolo, A., Kotler, L. A., Jarrett, M. A., & Spicer, J. (2007). Assessment and prevention of head motion during imaging of patients with attention deficit hyperactivity disorder. Psychiatry Research - Neuroimaging, 155(1), 75–82. https://doi.org/10.1016/j.pscychresns.2006.12.009

Fischl, B., & Dale, A. M. (2000). Measuring the thickness of the human cerebral cortex from magnetic resonance images. Proceedings of the National Academy of Sciences of the United States of America, 97(20), 11050–11055. https://doi.org/10.1073/pnas.200033797

Fischl, B., Sereno, M. I., & Dale, A. M. (1999). Cortical surface-based analysis: II. Inflation, flattening, and a surface-based coordinate system. NeuroImage, 9(2), 195–207. https://doi.org/10.1006/nimg.1998.0396

Fischl, B., Sereno, M. I., Tootell, R. B. H., & Dale, A. M. (1999). High-resolution intersubject averaging and a coordinate system for the cortical surface. Human Brain Mapping, 8(4), 272–284. https://doi.org/10.1002/(SICI)1097-0193(1999)8:4<272::AID-HBM10>3.0.CO;2-4

Fonov, V., Evans, A. C., Botteron, K., Almli, C. R., McKinstry, R. C., & Collins, D. L. (2011). Unbiased average age-appropriate atlases for pediatric studies. NeuroImage, 54(1), 313–327. https://doi.org/10.1016/j.neuroimage.2010.07.033

Garnett, E. O., Chow, H. M., Nieto-Castañón, A., Tourville, J. A., Guenther, F. H., & Chang, S.-E. (2018). Anomalous morphology in left hemisphere motor and premotor cortex of children who stutter. Brain, 141(9), 2670–2684. https://doi.org/10.1093/brain/awy199

Ghosh, S. S., Kakunoori, S., Augustinack, J., Nieto-Castanon, A., Kovelman, I., Gaab, N., Christodoulou, J. A., Triantafyllou, C., Gabrieli, J. D. E., & Fischl, B. (2010). Evaluating the validity of volume-based and surface-based brain image registration for developmental cognitive neuroscience studies in children 4 to 11 years of age. NeuroImage, 53(1), 85–93. https://doi.org/10.1016/j.neuroimage.2010.05.075

Greene, D. J., Black, K. J., & Schlaggar, B. L. (2016). Considerations for MRI study design and implementation in pediatric and clinical populations. Developmental Cognitive Neuroscience, 18, 101–112. https://doi.org/10.1016/j.dcn.2015.12.005

Guenette, J. P., Stern, R. A., Tripodis, Y., Chua, A. S., Schultz, V., Sydnor, V. J., Somes, N., Karmacharya, S., Lepage, C., Wrobel, P., Alosco, M. L., Martin, B. M., Chaisson, C. E., Coleman, M. J., Lin, A. P., Pasternak, O., Makris, N., Shenton, M. E., & Koerte, I. K. (2018). Automated versus manual segmentation of brain region volumes in former football players. NeuroImage: Clinical, 18, 888–896. https://doi.org/10.1016/j.nicl.2018.03.026

Hoogman, M., Muetzel, R., Guimaraes, J. P., Shumskaya, E., Mennes, M., Zwiers, M. P., Jahanshad, N., Sudre, G., Wolfers, T., Earl, E. A., Soliva Vila, J. C., Vives-Gilabert, Y., Khadka, S., Novotny, S. E., Hartman, C. A., Heslenfeld, D. J., Schweren, L. J. S., Ambrosino, S., Oranje, B.,… Franke, B. (2019). Brain imaging of the cortex in ADHD: A coordinated analysis of large-scale clinical and population-based samples. American Journal of Psychiatry, 176(7), 531–542. https://doi.org/10.1176/appi.ajp.2019.18091033

Kamson, D. O., Pilli, V. K., Asano, E., Jeong, J. W., Sood, S., Juhász, C., & Chugani, H. T. (2016). Cortical thickness asymmetries and surgical outcome in neocortical epilepsy. Journal of the Neurological Sciences, 368, 97–103. https://doi.org/10.1016/j.jns.2016.06.065

Karlsson, L., Tolvanen, M., Scheinin, N. M., Uusitupa, H.-M., Korja, R., Ekholm, E., Tuulari, J. J., Pajulo, M., Huotilainen, M., Paunio, T., & Karlsson, H. (2018). Cohort Profile: The FinnBrain Birth Cohort Study (FinnBrain). International Journal of Epidemiology, 47(1), 15–16j. https://doi.org/10.1093/ije/dyx173

Klapwijk, E. T., van de Kamp, F., van der Meulen, M., Peters, S., & Wierenga, L. M. (2019). Qoala-T: A supervised-learning tool for quality control of FreeSurfer segmented MRI data. NeuroImage, 189, 116–129. https://doi.org/10.1016/j.neuroimage.2019.01.014

Kuklisova-Murgasova, M., Aljabar, P., Srinivasan, L., Counsell, S. J., Doria, V., Serag, A., Gousias, I. S., Boardman, J. P., Rutherford, M. A., Edwards, A. D., Hajnal, J. V., & Rueckert, D. (2011). A dynamic 4D probabilistic atlas of the developing brain. NeuroImage, 54(4), 2750–2763. https://doi.org/10.1016/j.neuroimage.2010.10.019

Kumpulainen, V., Lehtola, S. J., Tuulari, J. J., Silver, E., Copeland, A., Korja, R., Karlsson, H., Karlsson, L., Merisaari, H., Parkkola, R., Saunavaara, J., Lähdesmäki, T., & Scheinin, N. M. (2020). Prevalence and Risk Factors of Incidental Findings in Brain MRIs of Healthy Neonates—The FinnBrain Birth Cohort Study. Frontiers in Neurology, 10, 1347. https://doi.org/10.3389/fneur.2019.01347

Kuperberg, G. R., Broome, M. R., McGuire, P. K., David, A. S., Eddy, M., Ozawa, F., Goff, D., West, W. C., Williams, S. C. R., Van der Kouwe, A. J. W., Salat, D. H., Dale, A. M., & Fischl, B. (2003). Regionally localized thinning of the cerebral cortex in schizophrenia. Archives of General Psychiatry, 60(9), 878–888. https://doi.org/10.1001/archpsyc.60.9.878

Lee, Y. J., Yum, M. S., Kim, M. J., Shim, W. H., Yoon, H. M., Yoo, I. H., Lee, J., Lim, B. C., Kim, K. J., & Ko, T. S. (2017). Large-scale structural alteration of brain in epileptic children with SCN1A mutation. NeuroImage: Clinical, 15, 594–600. https://doi.org/10.1016/j.nicl.2017.06.002

Lidauer, K., Pulli, E. P., Copeland, A., Silver, E., Kumpulainen, V., Merisaari, H., Saunavaara, J., Parkkola, R., Lähdesmäki, T., Saukko, E., Nolvi, S., Kataja, E., Korja, R., Karlsson, L., Karlsson, H., & Tuulari, J. J. (n.d.). Subcortical brain segmentation in 5-year-old children: validation of FSL-FIRST and FreeSurfer against manual segmentation.

Lyall, A. E., Shi, F., Geng, X., Woolson, S., Li, G., Wang, L., Hamer, R. M., Shen, D., & Gilmore, J. H. (2015). Dynamic Development of Regional Cortical Thickness and Surface Area in Early Childhood. Cerebral Cortex, 25(8), 2204–2212. https://doi.org/10.1093/cercor/bhu027

Masouleh, S. K., Eickhoff, S. B., Zeighami, Y., Lewis, L. B., Dahnke, R., Gaser, C., Chouinard-Decorte, F., Lepage, C., Scholtens, L. H., Hoffstaedter, F., Glahn, D. C., Blangero, J., Evans, A. C., Genon, S., & Valk, S. L. (2020). Influence of processing pipeline on cortical thickness measurement. Cerebral Cortex, 30(9), 5014–5027. https://doi.org/10.1093/cercor/bhaa097

McCarthy, C. S., Ramprashad, A., Thompson, C., Botti, J. A., Coman, I. L., & Kates, W. R. (2015). A comparison of FreeSurfer-generated data with and without manual intervention. Frontiers in Neuroscience, 9(OCT), 379. https://doi.org/10.3389/fnins.2015.00379

Merisaari, H., Tuulari, J. J., Karlsson, L., Scheinin, N. M., Parkkola, R., Saunavaara, J., Lähdesmäki, T., Lehtola, S. J., Keskinen, M., Lewis, J. D., Evans, A. C., & Karlsson, H. (2019). Test-retest reliability of Diffusion Tensor Imaging metrics in neonates. NeuroImage, 197, 598–607. https://doi.org/10.1016/j.neuroimage.2019.04.067

Morales, D. R., Slattery, J., Evans, S., & Kurz, X. (2018). Antidepressant use during pregnancy and risk of autism spectrum disorder and attention deficit hyperactivity disorder: Systematic review of observational studies and methodological considerations. BMC Medicine, 16(1), 6. https://doi.org/10.1186/s12916-017-0993-3

Muzik, O., Chugani, D. C., Juhász, C., Shen, C., & Chugani, H. T. (2000). Statistical parametric mapping: Assessment of application in children. NeuroImage, 12(5), 538–549. https://doi.org/10.1006/nimg.2000.0651

Nwosu, E. C., Robertson, F. C., Holmes, M. J., Cotton, M. F., Dobbels, E., Little, F., Laughton, B., van der Kouwe, A., & Meintjes, E. M. (2018). Altered brain morphometry in 7-year old HIV-infected children on early ART. Metabolic Brain Disease, 33(2), 523–535. https://doi.org/10.1007/s11011-017-0162-6

Oishi, K., Mori, S., Donohue, P. K., Ernst, T., Anderson, L., Buchthal, S., Faria, A., Jiang, H., Li, X., Miller, M. I., van Zijl, P. C. M., & Chang, L. (2011). Multi-contrast human neonatal brain atlas: Application to normal neonate development analysis. NeuroImage, 56(1), 8–20. https://doi.org/10.1016/j.neuroimage.2011.01.051

Phan, T. V., Sima, D. M., Beelen, C., Vanderauwera, J., Smeets, D., & Vandermosten, M. (2018). Evaluation of methods for volumetric analysis of pediatric brain data: The childmetrix pipeline versus adult-based approaches. NeuroImage: Clinical, 19, 734–744. https://doi.org/10.1016/j.nicl.2018.05.030

Phan, T. V., Smeets, D., Talcott, J. B., & Vandermosten, M. (2018). Processing of structural neuroimaging data in young children: Bridging the gap between current practice and state-of-the-art methods. In Developmental Cognitive Neuroscience (Vol. 33, pp. 206–223). Elsevier Ltd. https://doi.org/10.1016/j.dcn.2017.08.009

Poldrack, R. A., Paré-Blagoev, E. J., & Grant, P. E. (2002). Pediatric functional magnetic resonance imaging: Progress and challenges. Topics in Magnetic Resonance Imaging, 13(1), 61–70. https://doi.org/10.1097/00002142-200202000-00005

Pulli, E. P., Kumpulainen, V., Kasurinen, J. H., Korja, R., Merisaari, H., Karlsson, L., Parkkola, R., Saunavaara, J., Lähdesmäki, T., Scheinin, N. M., Karlsson, H., & Tuulari, J. J. (2019). Prenatal exposures and infant brain: Review of magnetic resonance imaging studies and a population description analysis. Human Brain Mapping, 40(6), 1987–2000. https://doi.org/10.1002/hbm.24480

Ranger, M., Chau, C. M. Y., Garg, A., Woodward, T. S., Beg, M. F., Bjornson, B., Poskitt, K., Fitzpatrick, K., Synnes, A. R., Miller, S. P., & Grunau, R. E. (2013). Neonatal Pain-Related Stress Predicts Cortical Thickness at Age 7 Years in Children Born Very Preterm. PLoS ONE, 8(10), e76702. https://doi.org/10.1371/journal.pone.0076702

Rodriguez, A., Miettunen, J., Henriksen, T. B., Olsen, J., Obel, C., Taanila, A., Ebeling, H., Linnet, K. M., Moilanen, I., & Järvelin, M. R. (2008). Maternal adiposity prior to pregnancy is associated with ADHD symptoms in offspring: Evidence from three prospective pregnancy cohorts. International Journal of Obesity, 32(3), 550–557. https://doi.org/10.1038/sj.ijo.0803741

Rodriguez, Alina. (2010). Maternal pre-pregnancy obesity and risk for inattention and negative emotionality in children. Journal of Child Psychology and Psychiatry, 51(2), 134–143. https://doi.org/10.1111/j.1469-7610.2009.02133.x

Roos, A., Jones, G., Howells, F. M., Stein, D. J., & Donald, K. A. (2014). Structural brain changes in prenatal methamphetamine-exposed children. Metabolic Brain Disease, 29(2), 341–349. https://doi.org/10.1007/s11011-014-9500-0

Rosas, H. D., Liu, A. K., Hersch, S., Glessner, M., Ferrante, R. J., Salat, D. H., Van Der Kouwe, A., Jenkins, B. G., Dale, A. M., & Fischl, B. (2002). Regional and progressive thinning of the cortical ribbon in Huntington’s disease. Neurology, 58(5), 695–701. https://doi.org/10.1212/WNL.58.5.695

Salat, D. H. (2004). Thinning of the Cerebral Cortex in Aging. Cerebral Cortex, 14(7), 721–730. https://doi.org/10.1093/cercor/bhh032

Schoemaker, D., Buss, C., Head, K., Sandman, C. A., Davis, E. P., Chakravarty, M. M., Gauthier, S., & Pruessner, J. C. (2016). Hippocampus and amygdala volumes from magnetic resonance images in children: Assessing accuracy of FreeSurfer and FSL against manual segmentation. NeuroImage, 129, 1–14. https://doi.org/10.1016/j.neuroimage.2016.01.038

Ségonne, F., Dale, A. M., Busa, E., Glessner, M., Salat, D., Hahn, H. K., & Fischl, B. (2004). A hybrid approach to the skull stripping problem in MRI. NeuroImage, 22(3), 1060–1075. https://doi.org/10.1016/j.neuroimage.2004.03.032

Seiger, R., Ganger, S., Kranz, G. S., Hahn, A., & Lanzenberger, R. (2018). Cortical Thickness Estimations of FreeSurfer and the CAT12 Toolbox in Patients with Alzheimer’s Disease and Healthy Controls. Journal of Neuroimaging, 28(5), 515–523. https://doi.org/10.1111/jon.12521

Serag, A., Kyriakopoulou, V., Rutherford, M. A., Edwards, A. D., Hajnal, J. V, Aljabar, P., Counsell, S. J., Boardman, J., & Rueckert, D. (2012). A Multi-channel 4D Probabilistic Atlas of the Developing Brain: Application to Fetuses and Neonates. Annals of the BMVA, 2012(3), 1–14. https://www.research.ed.ac.uk/en/publications/a-multi-channel-4d-probabilistic-atlas-of-the-developing-brain-ap

Shaw, P., Eckstrand, K., Sharp, W., Blumenthal, J., Lerch, J. P., Greenstein, D., Clasen, L., Evans, A., Giedd, J., & Rapoport, J. L. (2007). Attention-deficit/hyperactivity disorder is characterized by a delay in cortical maturation. Proceedings of the National Academy of Sciences of the United States of America, 104(49), 19649–19654. https://doi.org/10.1073/pnas.0707741104

Shi, F., Yap, P. T., Wu, G., Jia, H., Gilmore, J. H., Lin, W., & Shen, D. (2011). Infant brain atlases from neonates to 1- and 2-year-olds. PLoS ONE, 6(4). https://doi.org/10.1371/journal.pone.0018746

Sled, J. G., Zijdenbos, A. P., & Evans, A. C. (1998). A nonparametric method for automatic correction of intensity nonuniformity in mri data. IEEE Transactions on Medical Imaging, 17(1), 87–97. https://doi.org/10.1109/42.668698

Tanda, R., & Salsberry, P. J. (2014). Racial Differences in the Association Between Maternal Prepregnancy Obesity and Children’s Behavior Problems. Journal of Developmental & Behavioral Pediatrics, 35(2), 118–127. https://doi.org/10.1097/DBP.0000000000000007

Theys, C., Wouters, J., & Ghesquière, P. (2014). Diffusion tensor imaging and resting-state functional MRI-scanning in 5- and 6-year-old children: Training protocol and motion assessment. PLoS ONE, 9(4). https://doi.org/10.1371/journal.pone.0094019

Thompson, P. M., Jahanshad, N., Ching, C. R. K., Salminen, L. E., Thomopoulos, S. I., Bright, J., Baune, B. T., Bertolín, S., Bralten, J., Bruin, W. B., Bülow, R., Chen, J., Chye, Y., Dannlowski, U., de Kovel, C. G. F., Donohoe, G., Eyler, L. T., Faraone, S. V., Favre, P.,… Zelman, V. (2020). ENIGMA and global neuroscience: A decade of large-scale studies of the brain in health and disease across more than 40 countries. In Translational Psychiatry (Vol. 10, Issue 1, pp. 1–28). Springer Nature. https://doi.org/10.1038/s41398-020-0705-1

Vanderauwera, J., Altarelli, I., Vandermosten, M., De Vos, A., Wouters, J., & Ghesquière, P. (2018). Atypical Structural Asymmetry of the Planum Temporale is Related to Family History of Dyslexia. Cerebral Cortex, 28(1), 63–72. https://doi.org/10.1093/cercor/bhw348

Waters, A. B., Mace, R. A., Sawyer, K. S., & Gansler, D. A. (2019). Identifying errors in Freesurfer automated skull stripping and the incremental utility of manual intervention. Brain Imaging and Behavior, 13(5), 1281–1291. https://doi.org/10.1007/s11682-018-9951-8

Wedderburn, C. J., Subramoney, S., Yeung, S., Fouche, J. P., Joshi, S. H., Narr, K. L., Rehman, A. M., Roos, A., Ipser, J., Robertson, F. C., Groenewold, N. A., Gibb, D. M., Zar, H. J., Stein, D. J., & Donald, K. A. (2020). Neuroimaging young children and associations with neurocognitive development in a South African birth cohort study. NeuroImage, 219, 116846. https://doi.org/10.1016/j.neuroimage.2020.116846

White, T., Jansen, P. R., Muetzel, R. L., Sudre, G., El Marroun, H., Tiemeier, H., Qiu, A., Shaw, P., Michael, A. M., & Verhulst, F. C. (2018). Automated quality assessment of structural magnetic resonance images in children: Comparison with visual inspection and surface-based reconstruction. Human Brain Mapping, 39(3), 1218–1231. https://doi.org/10.1002/hbm.23911

Wilke, M., Holland, S. K., Altaye, M., & Gaser, C. (2008). Template-O-Matic: A toolbox for creating customized pediatric templates. NeuroImage, 41(3), 903–913. https://doi.org/10.1016/j.neuroimage.2008.02.056

Wilke, M., Schmithorst, V. J., & Holland, S. K. (2003). Normative pediatric brain data for spatial normalization and segmentation differs from standard adult data. Magnetic Resonance in Medicine, 50(4), 749–757. https://doi.org/10.1002/mrm.10606

Yang, D. Y. J., Beam, D., Pelphrey, K. A., Abdullahi, S., & Jou, R. J. (2016). Cortical morphological markers in children with autism: A structural magnetic resonance imaging study of thickness, area, volume, and gyrification. Molecular Autism, 7(1), 11. https://doi.org/10.1186/s13229-016-0076-x

Yang, X. R., Carrey, N., Bernier, D., & MacMaster, F. P. (2015). Cortical thickness in young treatment-naive children with ADHD. Journal of Attention Disorders, 19(11), 925–930. https://doi.org/10.1177/1087054712455501

Yoon, U., Fonov, V. S., Perusse, D., & Evans, A. C. (2009). The effect of template choice on morphometric analysis of pediatric brain data. NeuroImage, 45(3), 769–777. https://doi.org/10.1016/j.neuroimage.2008.12.046

Zöllei, L., Iglesias, J. E., Ou, Y., Grant, P. E., & Fischl, B. (2020). Infant FreeSurfer: An automated segmentation and surface extraction pipeline for T1-weighted neuroimaging data of infants 0–2 years. NeuroImage, 218, 116946. https://doi.org/10.1016/j.neuroimage.2020.116946

